# Macroevolutionary analysis of swallows revived the sight-line hypothesis

**DOI:** 10.1101/2022.10.03.510686

**Authors:** Masaru Hasegawa

## Abstract

The evolution of conspicuous ornamentation is often thought to be the consequence of sexual selection, but this might not always be the case. One such candidate is contrasting pale-dark facial color patterns in front of eyes in insectivorous birds and other animals (lore-forehead borderlines, hereafter). A sight-line hypothesis suggested that the contrasting color between lore and forehead assists in tracking and capturing a fast-moving prey. However, this classic hypothesis have been criticized (and thus ignored) for several reasons including lack of formal statistical test controlling for phylogenetic inertia and confounding effect of dark facial color markings that are beneficial by reducing glare. Here, using a phylogenetic comparative approach, we tested the sight-line hypothesis and a widespread alternative explanation, the sexual selection hypothesis, in hirundines (Aves: Hirundinidae). We found no support for the sexual function of lore-forehead borderline in hirundines, because lore-forehead borderline was not positively related to indices of sexual selection (sexual plumage dimorphism and extrapair mating opportunity). In contrast, we found consistent support for the sight-line hypothesis. Species foraging on large prey items (i.e., fast prey) had higher degrees of lore-forehead borderline than others in this clade. Furthermore, an analysis of evolutionary pathways suggested inter-dependent evolution of lore-forehead borderline and prey size; transitions to the state with large prey and no lore-forehead borderline were less likely to occur than transitions from that state. These results remained significant when excluding species that lack dark lore, and thus, not mere presence of dark lores, but contrasting color patterns would be important. To my knowledge, the current study is the first macroevolutionary support for the sight-line hypothesis.

## 1. Introduction

Many animals possess conspicuous traits, i.e., ornamentation, which seems to have rather negative effects on survival [1]. The evolution and maintenance of these traits are often explained by sexual selection, simply because of intuitive importance of ornamentation in mate attraction and rival deterrence (reviewed in [1,2]). However, this is clearly not the sole explanation. As is the case for aposematic coloration, conspicuous ornaments could have evolved for the positive effects on survival even if it is counterintuitive at first glance (e.g. see [3] for for poison frogs; also see [4] for the reviews of bird plumage coloration).

Facial color patterns such as forehead patches and eyebrow stripes might be another example. These traits were often found in insectivorous birds, such as flycatchers (e.g., the collared flycatcher, *Ficedula albicollis*) and swallows (e.g., the barn swallow, *Hirundo rustica*: Figs. 1 & S1; [5]). Although sexual selection on these traits has been reported (e.g., forehead patch size in flycatchers; [6]), the presence of sexual selection does not validate the hypothesis that sexual selection is responsible for the evolution and maintenance of the focal traits (sexual selection hypothesis, hereafter). In fact, contrasting pale feathers above dark feathers in front of eyes (lore-forehead borderline, hereafter) might be adaptive as a “sight line”, which can aid in tracking and capturing a fast-moving prey [5,7]. Then, the sexual function of lore-forehead borderline might be a byproduct and have only a limited effect on the evolution (i.e., exaptation: [8]). However, the sight-line hypothesis has never been tested in modern micro- and macroevolutionary approaches (and thus this kind of coloration is the least tested of all color patterns; [3]). Furthermore, previous support for this hypothesis is provided by simple interspecific comparison with a broad prey category (e.g., plants as well as animal preys) with no consideration of phylogenetic inertia [3,5,7], and thus, it is unclear whether or not contrasting “eyelines” co-evolved with “tracking and capturing fast-moving prey” (see above). In addition, previous studies of eyelines have been criticized, because their results can be explained solely by dark lore coloration, because dark lores can be regarded as a form of dark facial feather markings that are beneficial by reducing glare [3,9,10]. A macroevolutionary test using the same prey category with different moving speeds (e.g., various aerial insects), while controlling for phylogenetic inertia and dark lore coloration, is required.

**Figure 1.**
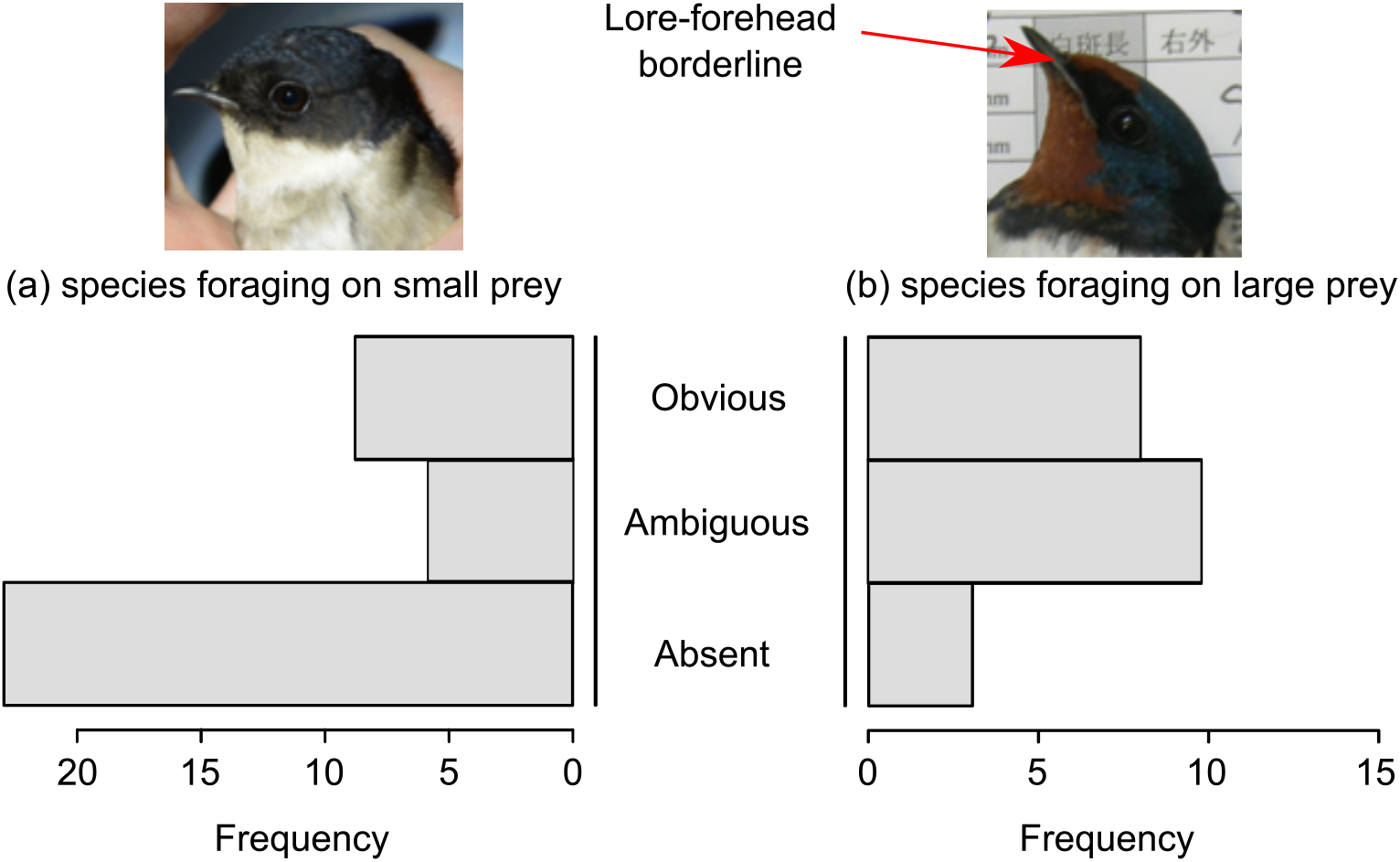
Histograms of species showing absent, ambiguous, and obvious lore-forehead borderline in (a) species foraging on small prey and (b) species foraging on large prey. Pictures above histograms are examples of species that forage on small prey (the Asian house martin, *Delichon dasypus*) and species that typically nest on buildings (the barn swallow *Hirundo rustica*). See text for a detailed explanation of each category of birds. Also see Fig. S7 for a closeup photograph of the faces of the two representative species.

Swallows and martins (family: Hirundinidae) are suitable systems for a macroevolutionary test of the sight-line hypothesis. All species are diurnal hyper-aerial insectivores [11,12], and hence the potential confounding effect of differential ecological background would be negligible. Unlike swifts (family: Apodidae), i.e., another hyper-aerial insectivorous bird, swallows have strong jaw muscles and thus can forage on large prey items [11,13], providing a good opportunity to test the sight-line hypothesis, because prey size is tightly linked to their flight speed (but is not tightly linked to maneuverability; [14]). In addition, although hirundines are typically socially monogamous birds with biparental provisioning [11]), they show variable extrapair mating opportunities: hirundines with biparental incubation had little extrapair paternity compared with those with female-only incubation, perhaps due to tradeoff between male incubation effort and his extrapair mating effort ([15,16]; also see [17] for smaller sperm size in hirundines with biparental incubation, reinforcing this perspective). Thus, I can examine the relative importance of the sight-line hypothesis and the sexual selection hypothesis in this clade. Finally, these predictor variables (i.e., prey size and incubation type) have repeatedly evolved in hirundines [16,18]. This would also be the case in lore-forehead borderline (e.g., the barn swallow *Hirundo rustica*, the cave swallow *Petrochelidon fulva*, the banded martin *Riparia cincta*, but not the blue swallow *Hirundo atrocaerulea* and the Asian house martin *Delichon dasypus*, possess lore-forehead borderline; Figs. 1 & S2; see the Results section for detailed information). Therefore, I can examine replicated evolutionary gain/loss of lore-forehead borderline in relation to the sight-line hypothesis and the sexual selection hypothesis in this clade. Because most hirundiens share dark lores, I can test the sight-line hypothesis, after excluding a possible confounding effect of mere presence of dark lores.

Here, using a Bayesian phylogenetic comparative framework, I tested the sight-line hypothesis in hirundines (Aves: Hirundinidae). I predicted that, if lore-forehead borderline can aid tracking and capturing fast-moving prey (i.e., sight-line hypothesis), species foraging on large (and thus fast-moving) prey items should have lore-forehead borderline (see above). This pattern would persist even after excluding species that lack dark lores, if lore-forehead borderline (i.e., sight-line), rather than dark lore coloration, is a key factor. Because our previous study found a correlated evolution of prey size and social foraging, in which species foraging on small prey items tended to be social foragers [18], I also examined the possible confounding effects of social foraging. In addition, I also tested whether a widespread alternative explanation, the sexual selection hypothesis, explains the evolution of this trait. I predicted that, if lore-forehead borderline has some sexual functions, lore-forehead borderline would be found in species with intense sexual selection [18]. For this purpose, I focused on sexual plumage dimorphism as an index of sexual selection on plumage coloration [18,19] as well as incubation type, which is a proxy of extrapair mating opportunities in hirundines (see above). Lastly, based on these results, I also conducted an analysis of evolutionary pathways to clarify the transition patterns of lore-forehead borderline in relation to foraging mode and sexual selection in an evolutionary time scale.

## 2. Methods

### (a) Data collection

As in previous studies [16,18,20,21], information on wing length as a measure of body size, prey size, and migratory habit (migrants or not) was obtained from Turner and Rose (1994). Prey size was dichotomized into small and large prey, based on the description of [11], in which detailed information of prey is available, using the criteria given in the previous study [22]. Detailed information is given elsewhere [18,22]. Information on social foraging (solitary/social) was obtained from Johnson et al. [23].

The degree of lore-forehead borderline is based on plates of Turner and Rose [11]. I examined the presence of pale feathers above dark lore, which would be adaptive for foraging on fast prey items [7]. Thus, in addition to species with a forehead patch as in the barn swallows (Fig. 1), species with “eyebrows” stripe (e.g., the banded martin *Riparia cincta*, which shows supraloral stripe) and caps (e.g., the wire-tailed swallow *Hirundo smithii*) were also classified as having a lore-forehead borderline (Fig. S1). I classified all species into absence, ambiguous, and obvious lore-forehead borderline based on the sharpness of the borderline. I did not include iridescent shortwave (i.e., blue-green, or sometimes violet) coloration (e.g., see Fig. 1 for bluish dorsal coloration in the Asian house martin), because brightness (as well as hue and chroma) of such coloration depends on the angle to the light source, which would be unsuitable for birds to use as a sight line. Because plates of Turner and Rose [11] sometimes lack sex-separate plates, I also took into account additional literature [24] which showed sex-specific color patterns. In all three species that shows sex difference in degree of lore-forehead borderline, females, but not males, possess lore-forehead borderline (*Progne dominicensis, Progne modesta, Progne chalybea*: see Table S1). Because my working hypothesis, sight-line hypothesis, is concerning viability selection, I focused on female values, which would be closer to viability optima than males. However, I also reported results when I used male values, which would clarify the importance of sexual selection relative to sight-line hypothesis.

As a proxy of extra-pair mating opportunities (frequent/rare), I used incubation type (female-only versus biparental incubation) from a previous study [16], which were in turn obtained mainly from Turner and Rose [11]. An alternative approach using extrapair paternity itself instead of incubation type is impractical, because information of extrapair paternity is limited in this clade (only 10 species: [16]). In addition, I also recorded notable presence (or absence) of male-female overall plumage discrepancy, i.e., a measure of sexual plumage dimorphism, which can be collected from the notes for each plate of species in the text of Turner and Rose [11]. I included all kinds of sexual plumage dimorphism here such as tail length and body coloration as well as facial color pattern. The dataset is summarized in Table S1 (note that the scientific names follow birdtree.org; e.g., all genus “*Cecropis*” species are listed as genus “*Hirundo*”).

### (b) Statistics

I used a Bayesian phylogenetic mixed model with an ordinal error distribution to examine the degree of lore-forehead borderline. Due to limited sample size for predictor variables, I separately analyzed for each predictor together with confounding variables (i.e., body size and migratory habit). To account for phylogenetic uncertainty, I fit the models to each tree and applied multimodel inference using 1000 alternative trees for swallows from birdtree.org [25]. I derived mean coefficients, 95% credible intervals (CIs) based on the highest posterior density, and MCMC-based P-values (P_MCMC_), together with the phylogenetic signal [26]. All analyses were conducted in R ver. 4.0.0 [27] using the function “MCMCglmm” in the package “MCMCglmm” (ver. 2.29: [28]). I used a Gelman prior for the fixed effects while standardizing each continuous variable using “gelman.prior” in the package “MCMCglmm.” I ran the analysis for 140,000 iterations, with a burn-in period of 60,000 and a thinning interval of 80 for each tree.

I also used the discrete module in BayesTraits [29,30] to examine evolutionary transitions among states with lore-forehead borderline and foraging mode or sexual selection (as in [16] and other recent literatures in evolutionary ecology; e.g. [31]). Regarding foraging mode, I focused on large versus small prey size and social versus solitary foraging. Regarding sexual selection, I focused on sexual plumage dimorphism/monomorphism and incubation type, which is tightly linked to extra-pair paternity both between-species and within-species in this clade (see Introduction). I used 1000 alternative trees of Hirundininae from birdtree.org, which was sufficient to control for phylogenetic uncertainty [32]. Here, I ran for 1010000 iterations with a burn-in period of 10000 and a thinning interval of 1000 (i.e., I used MCMC methods in BayesTraits; note that a phylogenetic tree is chosen randomly from 1000 trees in each iteration). I denoted means as the representatives of each transition rate. Likewise, I showed mean differences in transition rates and the posterior probability that the differences would be higher (or lower) than zero (as P_MCMC_ values). The reproducibility of the Markov chain Monte Carlo (MCMC) simulation was confirmed by calculating the Brooks–Gelman–Rubin statistic (Rhat), which should be < 1.2 for all parameters [33], after repeating the analyses three times.

The Bayes factor (BF) was calculated by comparing the marginal likelihood of a dependent model that assumed correlated evolution of the focal two traits to that of an independent model assuming that the two traits evolved independently, using the stepping stone sampler implemented in BayesTraits (as suggested by BayesTraits Manual V.3). Bayes factors of >2, 5–10, and >10 indicate positive, strong, and very strong evidence of correlated evolution, respectively. Note that the range of transition rate is 10e-32 to 100 (not 0 to 1), as explained in BayesTraits Manual V.3. To visually check replicated co-distribution [34], I also presented an example of ancestral character reconstruction using the functions ‘ace’ in the R package ‘ape’ and ‘plotTree’ in the R package ‘phytools’ [35]. I confirmed that all statistical results supporting correlated evolution (see Results) did in fact represent replicated co-distributions throughout hirundines and thus false positives due to a few influential evolutionary events would be unlikely (as explained in other comparative studies: e.g., [36]). I also applied a threshold model [37], which is immune to within-clade pseudoreplication, using the function ‘threshBayes’ in the R package ‘phytools’

## 3. Results

### (a) Lore-forehead borderline across species

Lore-forehead borderlines are widely distributed in the family Hirundinidae through multiple evolutionary changes of the state (i.e., absent, ambiguous, and obvious lines; Figs. 2 & S2). When controlling for phylogeny and two covariates, log(wing length) as a measure of body size and migratory habit, species foraging on large prey items had a higher degree of lore-forehead borderline than those foraging on small prey items in female hirundines (Table 1a; Fig. 1). In fact, the interspecific distribution of foraging on large prey items closely matched the distribution of at least some degrees of lore-forehead borderline (Fig. 2). Similar, but non-significant, trends were found in males (Table 1a). When I used another measure of foraging mode, social foraging (solitary/social) instead of prey size, no detectable relationship was found in males and females (Table 1b; Fig. S3).

**Table 1.**
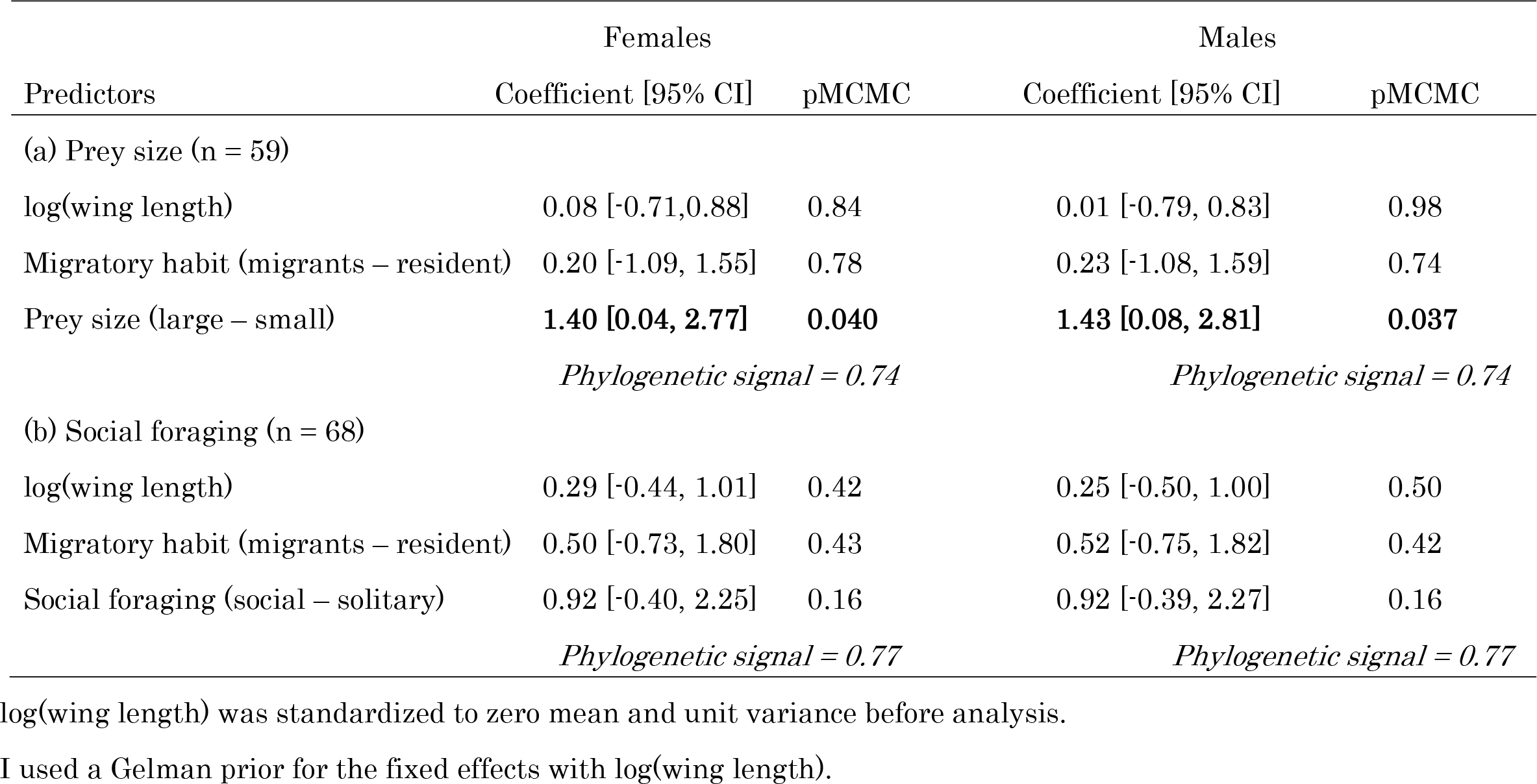
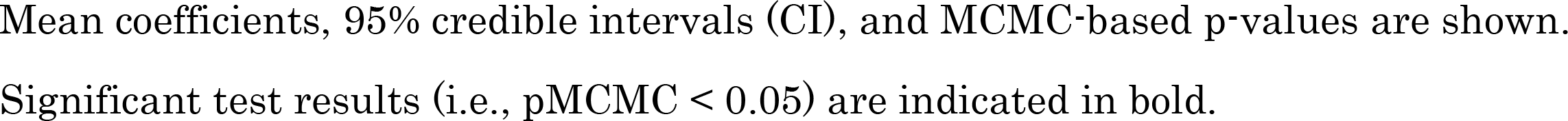
Multivariable Bayesian phylogenetic mixed model with an ordinal distribution predicting the degree of lore-forehead borderline (0: absent; 1: ambiguous; 2: obvious) in relation to foraging mode (a: prey size and b: social foraging) and covariates in female and male hirundines.

|  | Females |  | Males |  |
| --- | --- | --- | --- | --- |
| Predictors | Coefficient [95% CI] | pMCMC | Coefficient [95% CI] | pMCMC |
| (a) Prey size (n = 59) |  |  |  |  |
| log(wing length) | 0.08 [-0.71,0.88] | 0.84 | 0.01 [-0.79, 0.83] | 0.98 |
| Migratory habit (migrants – resident) | 0.20 [-1.09, 1.55] | 0.78 | 0.23 [-1.08, 1.59] | 0.74 |
| Prey size (large – small) | **1.40 [0.04, 2.77]** | **0.040** | **1.43 [0.08, 2.81]** | **0.037** |
|  | *Phylogenetic signal = 0.74* |  | *Phylogenetic signal = 0.74* |  |
| (b) Social foraging (n = 68) |  |  |  |  |
| log(wing length) | 0.29 [-0.44, 1.01] | 0.42 | 0.25 [-0.50, 1.00] | 0.50 |
| Migratory habit (migrants – resident) | 0.50 [-0.73, 1.80] | 0.43 | 0.52 [-0.75, 1.82] | 0.42 |
| Social foraging (social – solitary) | 0.92 [-0.40, 2.25] | 0.16 | 0.92 [-0.39, 2.27] | 0.16 |
|  | *Phylogenetic signal = 0.77* |  | *Phylogenetic signal = 0.77* |  |

log(wing length) was standardized to zero mean and unit variance before analysis.
I used a Gelman prior for the fixed effects with log(wing length).
Mean coefficients, 95% credible intervals (CI), and MCMC-based p-values are shown.
Significant test results (i.e., $p_{\text{MCMC}} < 0.05$ ) are indicated in bold.

**Figure 2.**
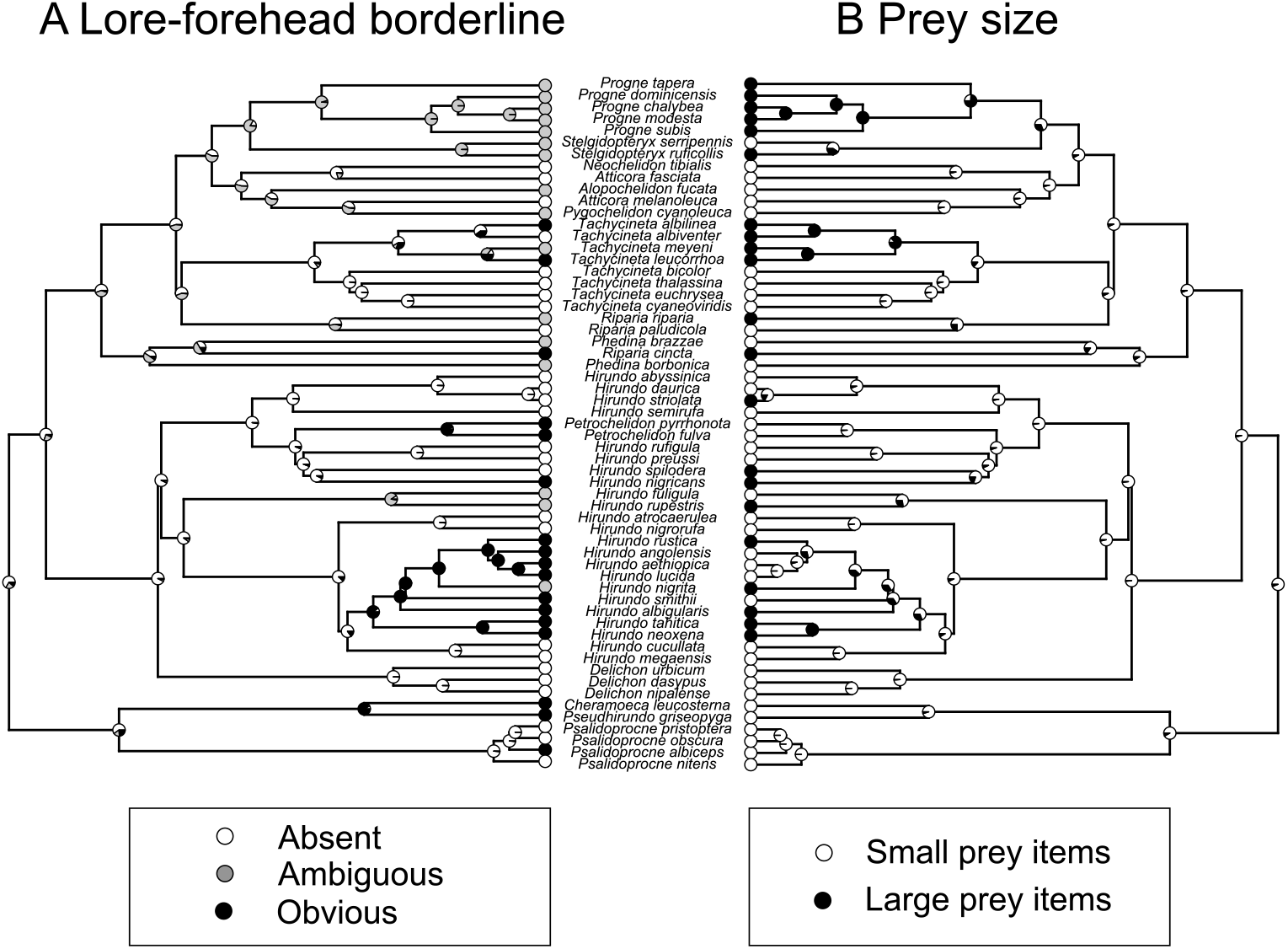
Examples of ancestral character reconstruction of the presence/absence of female lore-forehead borderline (left panel) and prey size (right panel) in swallows and martins (Aves: Hirundininae) using the functions ‘plotTree’ in the R package ‘phytools’ (Revell 2012). Black, grey, and white circles in tips in the left panel indicate species with obvious, ambiguous, and no lore-forehead borderline and black and white circles in tips in the right panel indicate species foraging on large and small prey, respectively. Likewise, the proportions of black, grey, and white in nodes in the left panel indicate the probability of an ancestral state with obvious, ambiguous, and no lore-forehead borderline, and corresponding black and white proportion in the right panel indicate the probability of an ancestral state foraging on large and small prey items.

When I used each of the two measures of sexual selection (i.e., sexual plumage dimorphism as a measure of sex-biased selection, or incubation type as a measure of opportunity for extrapair mating), neither variable showed significant relationships with the degree of lore-forehead borderline in male and female hirundines (Table 2; Figs. S2 & S4).

**Table 2.**
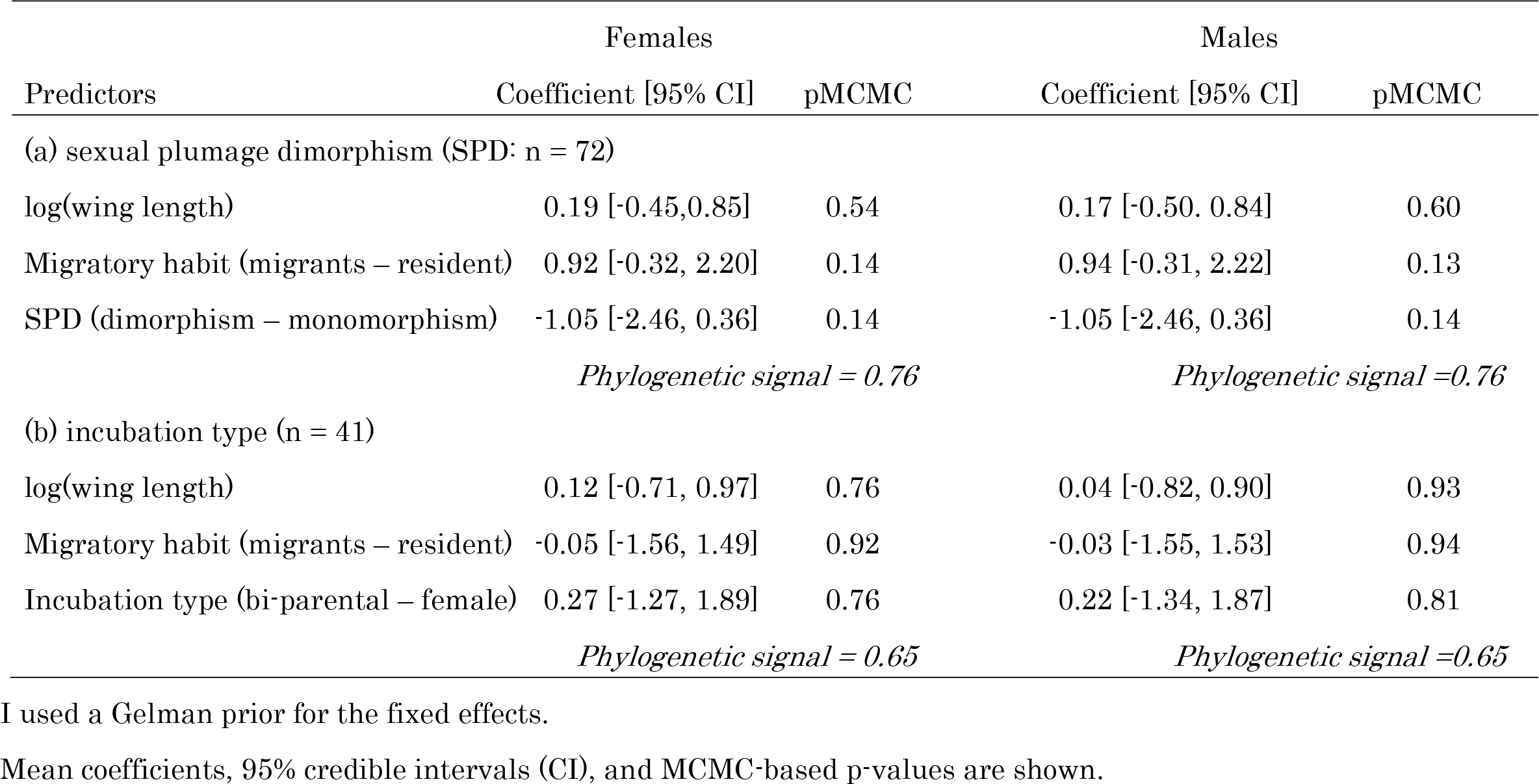
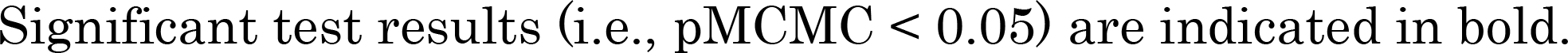
Multivariable Bayesian phylogenetic mixed model with an ordinal distribution predicting the degree of lore-forehead borderline (0: absent; 1: ambiguous; 2: obvious) in relation to measures of sexual selection (a: sexual plumage dimorphism and b: incubation type) and covariates in female and male hirundines.

|  | Females |  | Males |  |
| --- | --- | --- | --- | --- |
| Predictors | Coefficient [95% CI] | pMCMC | Coefficient [95% CI] | pMCMC |
| (a) sexual plumage dimorphism (SPD: n = 72) |  |  |  |  |
| log(wing length) | 0.19 [-0.45,0.85] | 0.54 | 0.17 [-0.50, 0.84] | 0.60 |
| Migratory habit (migrants – resident) | 0.92 [-0.32, 2.20] | 0.14 | 0.94 [-0.31, 2.22] | 0.13 |
| SPD (dimorphism – monomorphism) | -1.05 [-2.46, 0.36] | 0.14 | -1.05 [-2.46, 0.36] | 0.14 |
|  | *Phylogenetic signal = 0.76* |  | *Phylogenetic signal =0.76* |  |
| (b) incubation type (n = 41) |  |  |  |  |
| log(wing length) | 0.12 [-0.71, 0.97] | 0.76 | 0.04 [-0.82, 0.90] | 0.93 |
| Migratory habit (migrants – resident) | -0.05 [-1.56, 1.49] | 0.92 | -0.03 [-1.55, 1.53] | 0.94 |
| Incubation type (bi-parental – female) | 0.27 [-1.27, 1.89] | 0.76 | 0.22 [-1.34, 1.87] | 0.81 |
|  | *Phylogenetic signal = 0.65* |  | *Phylogenetic signal =0.65* |  |

I used a Gelman prior for the fixed effects.
Mean coefficients, 95% credible intervals (CI), and MCMC-based p-values are shown.
Significant test results (i.e., $p_{\text{MCMC}} < 0.05$ ) are indicated in bold.

The majority of species that lack a lore-forehead borderline has dark lores (females: 78%, 27/35; males: 79%, 30/38), and thus these effects would be mainly due to the presence/absence of lore-forehead borderline itself rather than the presence/absence of dark lore. In fact, when excluding the eight species that lack dark lores, qualitatively similar results were found in females (i.e., significant and nonsignificant remain unchanged; Tables S2 & S3). This was also the case in males, though this time prey size was marginal (P_MCMC_ = 0.053), whereas sexual dimorphism became significant, in which sexual dimorphism was negatively related to the degree of lore-forehead borderline (P_MCMC_ = 0.042; Tables S2 & S3).

In the eight species, in which both males and females lack dark lores, four species belong to the genus *Cecropis*, the remainings belong to the genus *Petrochelidon* (though the forest swallow, *P. fuliginosa*, was recently shown not to be *Petrochelidon*; [38]). This means that a limited number of evolutionary changes of lore coloration had occurred (min. one to max. three times evolution of light lore coloration), making it impossible to analyze the presence/absence of dark lore coloration.

### (b) Evolutionary pathway

Because the difference between species with large and small prey items is due largely to the presence/absence of lore-forehead borderline (Figs. 1 & 2), I examined an analysis of evolutionary pathways using two state variables, lore-forehead borderline (presence/absence) and prey size (large/small). I found strong support for correlated evolution of female lore-forehead borderline and prey size (n = 65, BF = 9.24; Fig. 3). Transition rate to the state with large prey items and no lore-forehead borderline was lower than the reverse transition (P_MCMC_ = 0.002), indicating that foraging on large prey items are less likely to evolve when hirundines lack lore-forehead borderline (note the multiple evolutionary events of evolutionary loss of lore-forehead borderline; Fig. 3). When using the threshold model, which would be less affected by false positive (see Methods), instead of BayesTraits, I found a similar positive correlation between the two binary traits (Fig. S5), again supporting the inter-dependent evolution. Qualitatively similar, but weaker, relationship was found when I used male values (BF = 3.87), though this yielded nonsignificant correlation when I applied a threshold model (r = 0.33, P_MCMC_ = 0.13).

**Figure 3.**
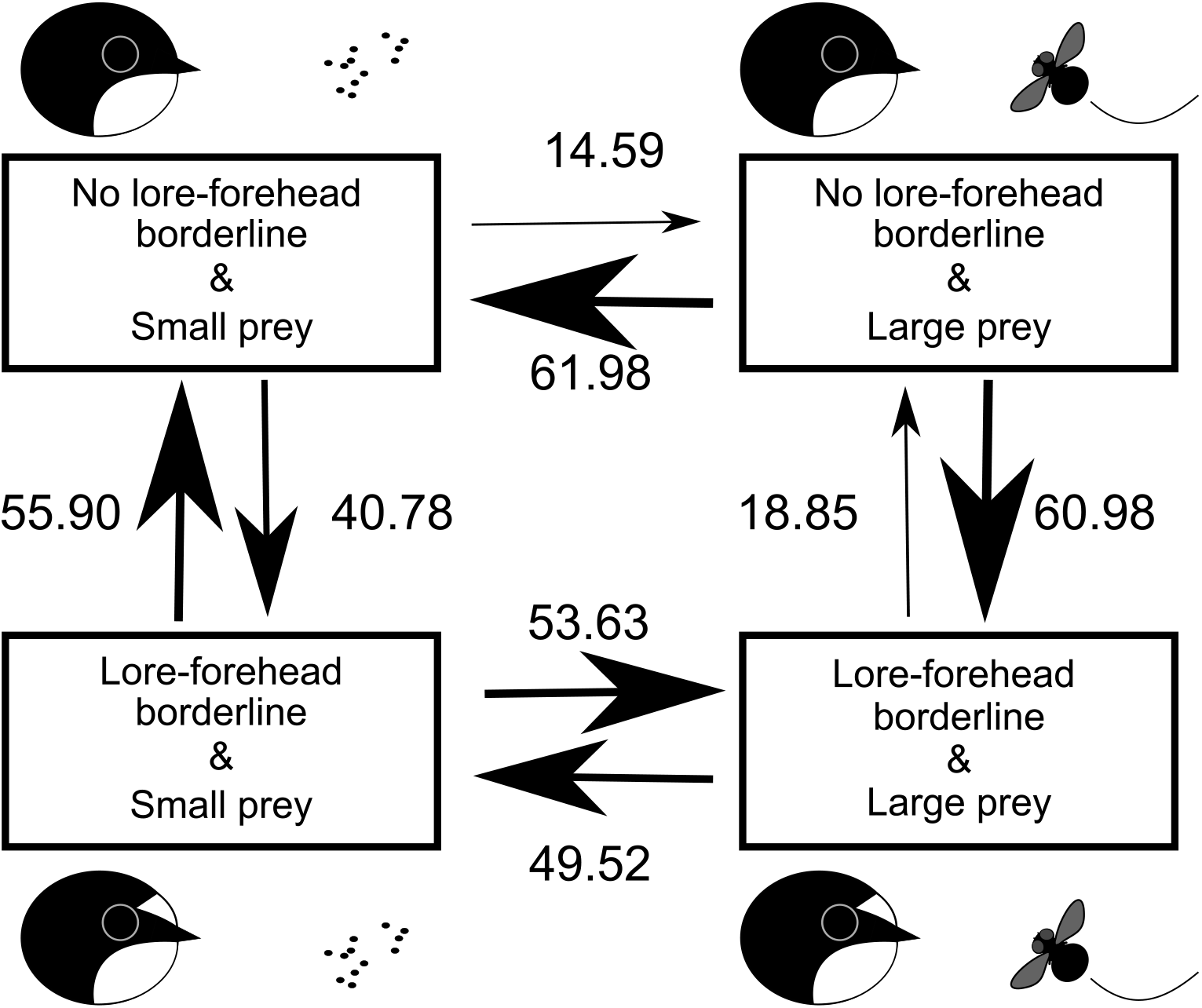
The most likely evolutionary transition between the presence and absence of lore-forehead borderline in relation to prey size (see text) in the family Hirundinidae. Model-averaged transition rates, which are reflected by the arrow size, are depicted. An illustration beside each state was added for easy understanding (also see Fig. S1 for detailed categorization of birds).

Similar analysis of evolutionary pathways using social foraging (solitary/social) and prey size yielded no detectable support for correlated evolution between the two state variables in both males and females (males: n = 68, BF = 0.02; females: n = 68, BF = -1.00; note that negative values can be estimated; see BayesTraits manual; www.evolution.rdg.ac.uk).

Although I found no detectable support for the correlated evolution between sexual plumage dimorphism and female lore-forehead borderlines (n = 72, BF = -0.06), I found positive support for correlated evolution of sexual plumage dimorphism and male lore-forehead borderline (n = 72, BF = 3.37). However, the observed pattern was opposite to the sexual selection explanation, because transition rates to the state with sexual plumage dimorphism and lore-forehead borderline was less likely to occur than the reverse transition (P_MCMC_ = 0.004; Fig. S6). When using the threshold model, the relationship between prey sexual plumage dimorphism and lore-forehead borderline was nonsignificant (r = -0.37, P_MCMC_ = 0.13), and thus the relationship was at best weak. When I used incubation type, i.e., another measure of sexual selection (see above), I found no detectable support for correlated evolution of lore-forehead borderline and incubation type (n = 41, BF = -0.12, 0.15 for males and females, respectively).

Qualitatively similar results were found when excluding eight species that lack dark lores (females: prey size: BF = 9.78; all others: BF < 0.54; males: prey size: BF = 3.79; sexual dimorphism: BF = 4.49; others: BF < -0.38). When using the threshold model, I found a positive correlation between prey size and lore-forehead borderline in females (r = 0.58, P_MCMC_ = 0.019), supporting the inter-dependent evolution. The relationship between prey size and lore-forehead borderline was nonsignificant in males when I applied a threshold model (r = 0.45, P_MCMC_ = 0.07). This was also the case when studying relationship between lore-forehead borderline and sexual plumage dimorphism (r = -0.36, P_MCMC_ = 0.12).

## 4. Discussion

The main finding of the current study is correlated evolution of lore-forehead borderline and prey size, in which evolutionary gain of foraging on large prey items are less likely to occur when hirundines lost lore-forehead borderline, supporting the sight-line hypothesis. Species that lack dark lore was rare in hirundines and excluding them from analyses did not change results qualitatively, and thus, not mere presence of dark lores, but contrasting pale-dark feather color patterns in front of eyes would be important. On the contrary, I could not find any support for the sexual selection hypothesis. Rather, sexual selection would weaken the evolutionary link between lore-forehead borderline and prey size through the weakly negative correlated evolution with lore-forehead borderline. Sexual selection would be a weak evolutionary force for lore-forehead borderline, which is also consistent with weak and fluctuating selection on the forehead patch in the pied flycatcher, a model species for sexual selection studies (reviewed in [39]; though I did not test the function of forehead patch itself). An alternative explanation that the current measures of sexual selection did not capture the interspecific variation in sexual selection is unlikely, because these measures were tightly linked to other sexual traits including the famous sexual trait, fork tails, in the same clade [16,18,20,21]. The current findings, therefore, support the sight-line hypothesis, i.e., the foraging function of lore-forehead borderline, rather than the sexual selection hypothesis.

Swallows and martins are hyper-aerial insectivores and thus it is straightforward that selection favors mechanisms promoting successful aerial foraging. In fact, they have a unique visual system with a narrow binocular field, bifoveate retina, and unusual long eye, which are similar to diurnal raptors such as hawks and falcons rather than other passerine birds [40]. Then, it is not surprising to see feather characteristics that enhance visual systems for successful aerial foraging. Recent studies have reevaluated dark facial feathers which reduce glare (e.g., [9,10]), but light-colored feathers in front of the eye clearly do not have this function and rather increase glare (see Results; note that Bortolotti [3] failed to take this into account and misinterpretted eye line as a form of dark facial feathers, whereas an ealier review pointed out the importance of contrasting coloration of eye line: [41]). As the classic sight-line hypothesis proposed [9,10], the contrasting feather coloration in front of the eye would still be adaptive to precisely aim their preys, and thus would be less likely to lost in hirundines foraging on large, speedy prey items. As the sighting device of fire arms increases successful shooting, lore-forehead borderline as a sight line would also increase foraging success when hirundines shoot out themselves toward fast-moving prey. To my knowledge, the current study is the first validation of the sight-line hypothesis in the modern macroevolutioanry approach, though direct behavioral evidence while manipulating lore-forehead borderlines is clearly needed.

Of course, this is not the sole explanation. Because the current study is a correlational study, I could not completely exclude the possibility that lore-forehead borderline might indirectly be related to prey size. For example, as is the case for “badge of status” (which in fact includes white forehead patch in flycatchers: reviewed in [39]), lore-forehead borderline of hirundines might have some functions to repel intra- and interspecific competitors for non-sexual resources including high-quality foods (i.e., large prey items; Turner 1982). In addition, as white tail spots can work as an illusion to increase the perceived length of outermost tail feathers in hirundines [21,42], lore-forehead borderline might have a similar psychological function on the perceived bill length, i.e., a dark-colored weapon of hirundines, providing competitive advantages (see [7] for a similar argument on facial marking in antelopes, which might increase the perceived length of the antlers; also see Fig. 3). These explanations are plausible at least in theory, but social foraging, which accompanies cooperative foraging for abundant prey items (i.e., limited competition; [18,43]), had no detectable relationship with lore-forehead borderline, indicating that the signaling function of lore-forehead borderline during foraging would be weak. No detectable sexual selection (including intra-sexual selection) favoring lore-forehead borderline reinforces this perspective (note that the opposite pattern was observed; see above). Therefore, these effects on the evolution of lore-forehead borderline via sexual (and social) selection would be at best small.

In summary, the current study supported the sight-line hypothesis, the foraging function (rather than sexual function) of lore-forehead borderline in hirundines. Researchers often ignore classic hypothesis while favoring sophisticated modern hypothesis. In fact, foraging on large prey is thought to be a selection force favoring deeply forked tails through its aerodynamic function (though this is logically invalid; reviewed in [18]), whereas virtually no studies focus on facial feathers though it clearly affects visions in these visual insectivores. Rather than targeting a highly-sophisticated, “eye-catching” hypothesis alone, we should take into account classic, but often overlooked, hypotheses as well, to understand the actual mechanisms of evolution and diversification of organisms.

## Ethics

This comparative study does not include any treatments of animals, as all the information was gathered from literatures.

## Data accessibility

The data sets supporting this article have been uploaded as part of electronic supplementary material, table S1, which will be uploaded to osf.io after acceptance.

## Authors’ contributions

I performed data analysis and wrote the manuscript alone.

## Acknowledgments

I thank Dr Emi Arai, Dr Shumpei Kitamura and his lab members at Ishikawa Prefectural University for their kindest advices. I am grateful to Dr Angela Turner for her kindly support on the valuable information on swallows.

## Figure legends

**Table S1.**
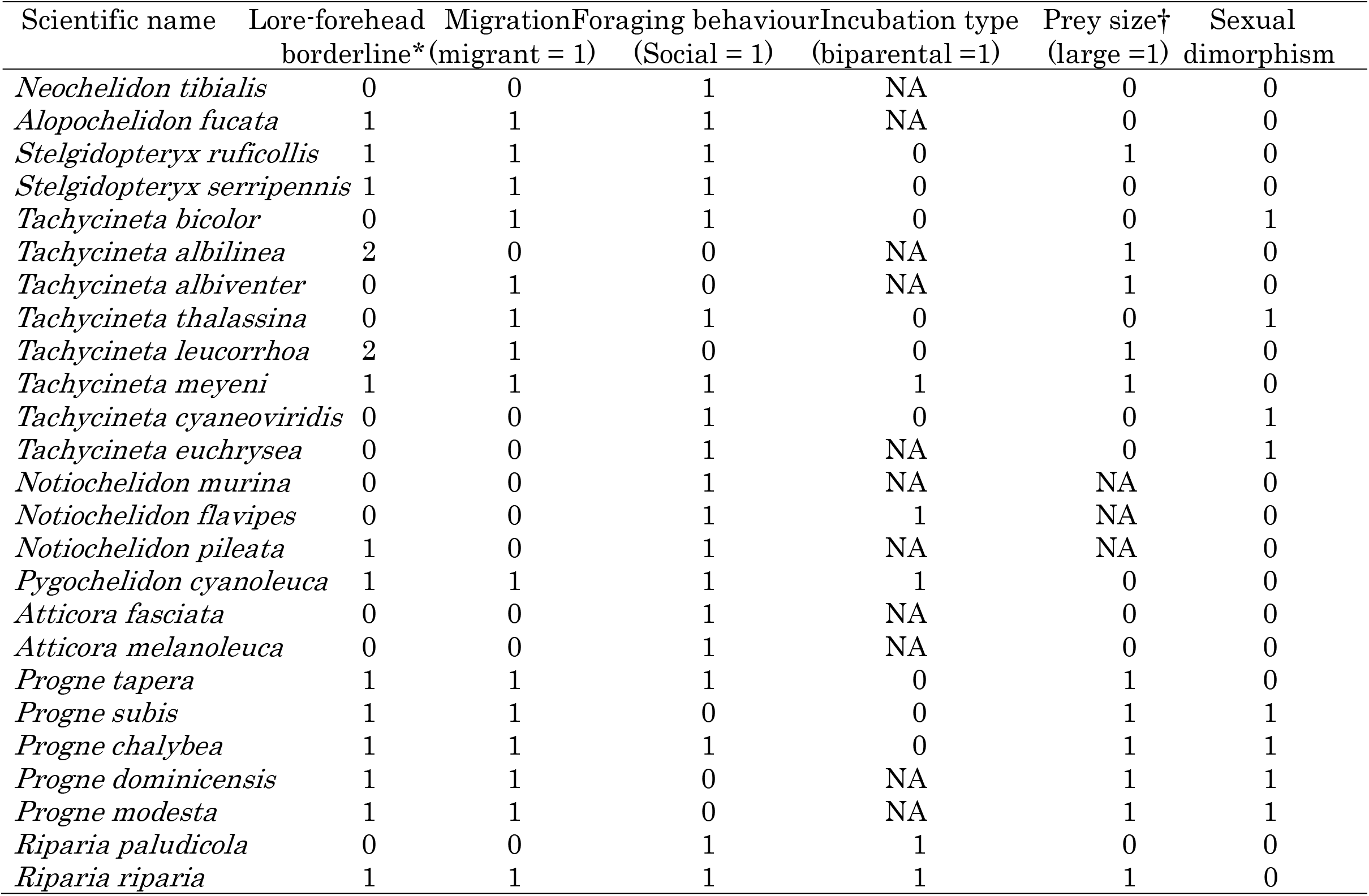
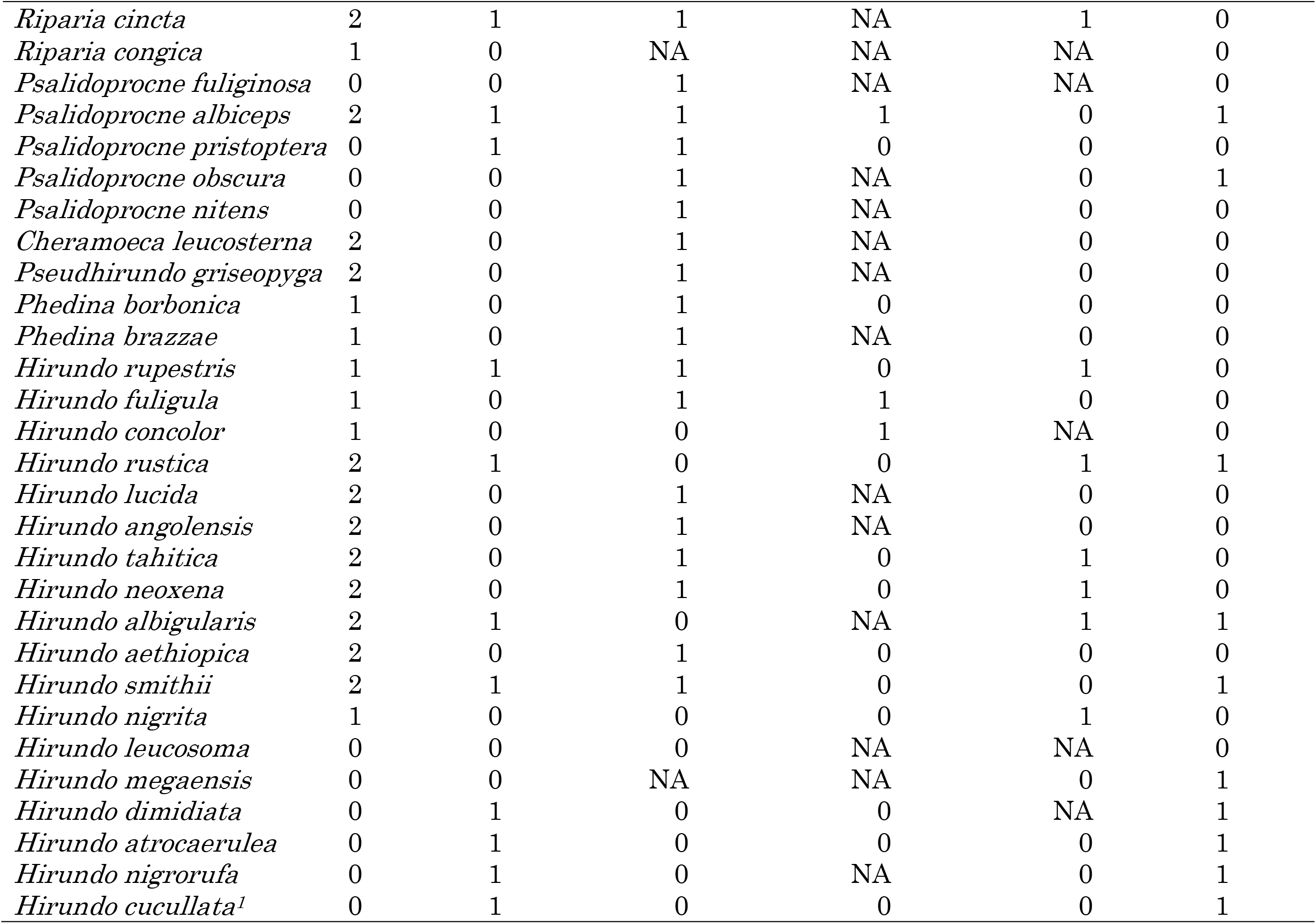
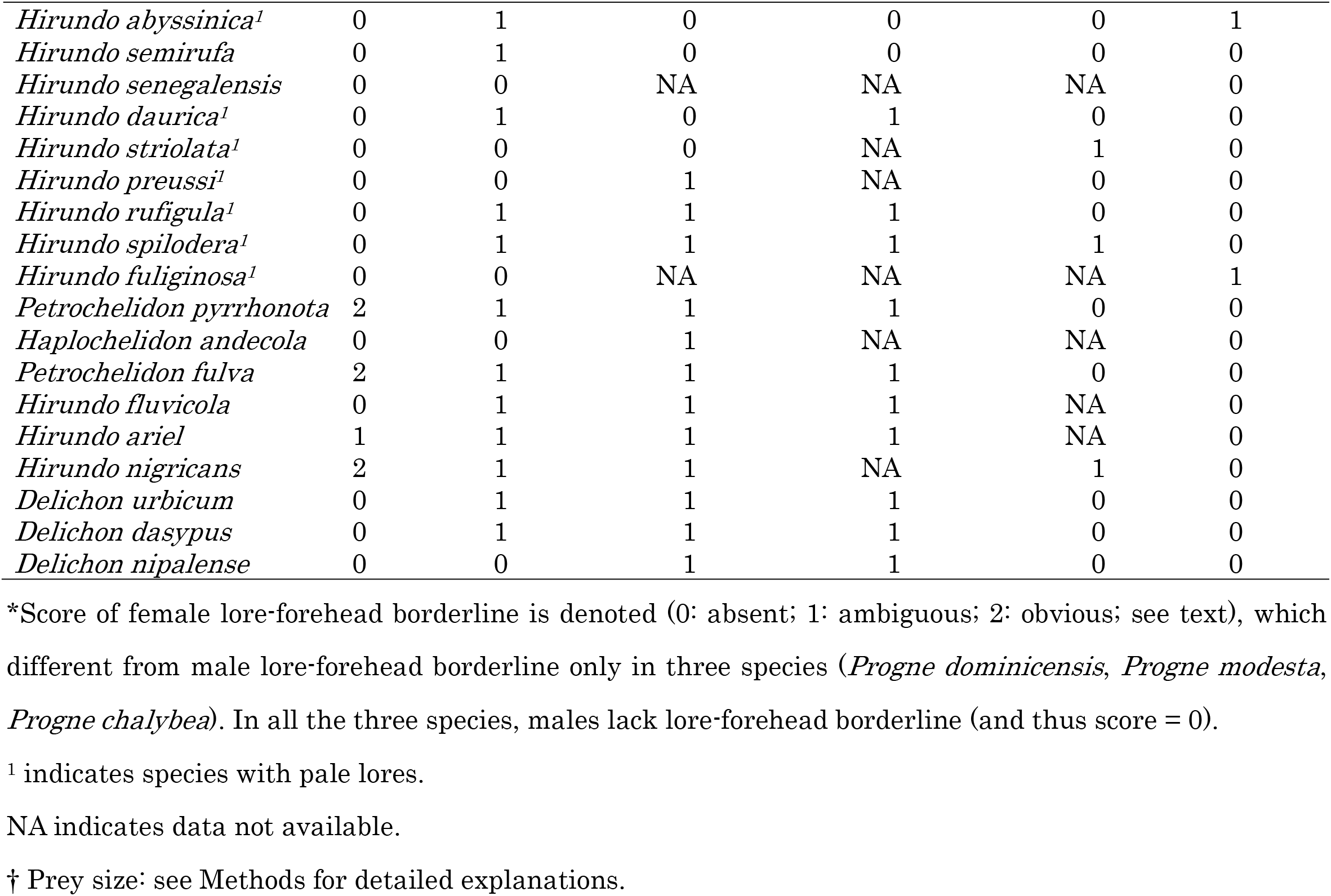
Data set for the current study in the subfamily Hirundininae (n = 72).

**Table S2.**
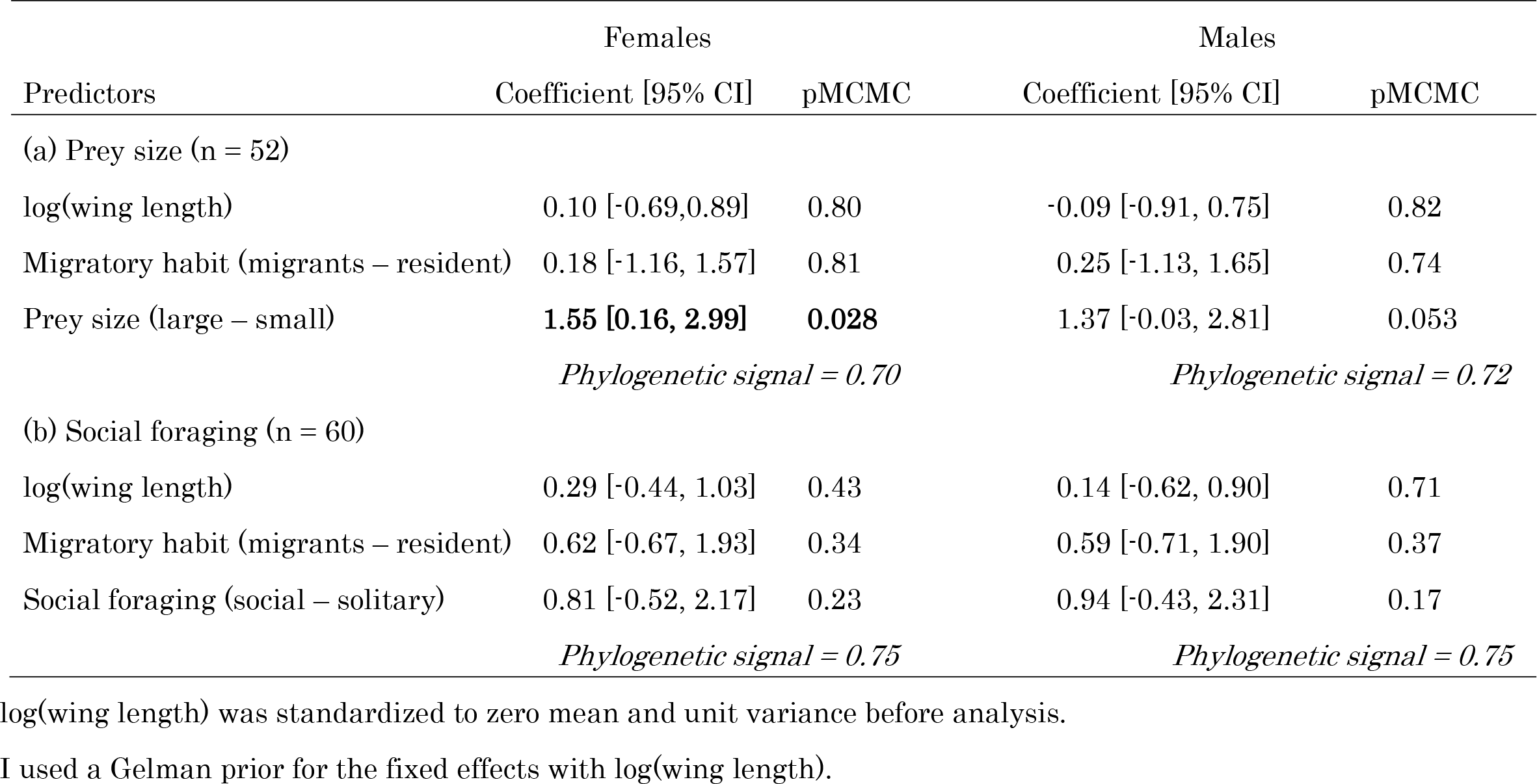
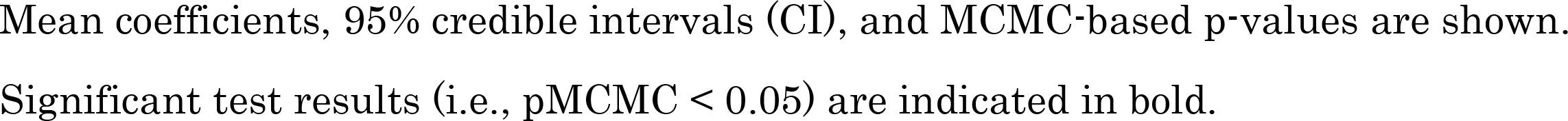
Multivariable Bayesian phylogenetic mixed model with an ordinal distribution predicting the degree of lore-forehead borderline (0: absent; 1: ambiguous; 2: obvious) in relation to foraging mode (a: prey size and b: social foraging) and covariates in female and male hirundines after excluding species with pale lores.

|  | Females |  | Males |  |
| --- | --- | --- | --- | --- |
| Predictors | Coefficient [95% CI] | pMCMC | Coefficient [95% CI] | pMCMC |
| (a) Prey size (n = 52) |  |  |  |  |
| log(wing length) | 0.10 [-0.69,0.89] | 0.80 | -0.09 [-0.91, 0.75] | 0.82 |
| Migratory habit (migrants – resident) | 0.18 [-1.16, 1.57] | 0.81 | 0.25 [-1.13, 1.65] | 0.74 |
| Prey size (large – small) | **1.55 [0.16, 2.99]** | **0.028** | 1.37 [-0.03, 2.81] | 0.053 |
|  | *Phylogenetic signal = 0.70* |  | *Phylogenetic signal = 0.72* |  |
| (b) Social foraging (n = 60) |  |  |  |  |
| log(wing length) | 0.29 [-0.44, 1.03] | 0.43 | 0.14 [-0.62, 0.90] | 0.71 |
| Migratory habit (migrants – resident) | 0.62 [-0.67, 1.93] | 0.34 | 0.59 [-0.71, 1.90] | 0.37 |
| Social foraging (social – solitary) | 0.81 [-0.52, 2.17] | 0.23 | 0.94 [-0.43, 2.31] | 0.17 |
|  | *Phylogenetic signal = 0.75* |  | *Phylogenetic signal = 0.75* |  |

log(wing length) was standardized to zero mean and unit variance before analysis.
I used a Gelman prior for the fixed effects with log(wing length).
Mean coefficients, 95% credible intervals (CI), and MCMC-based p-values are shown.
Significant test results (i.e., $p_{\text{MCMC}} < 0.05$ ) are indicated in bold.

**Table S3.**
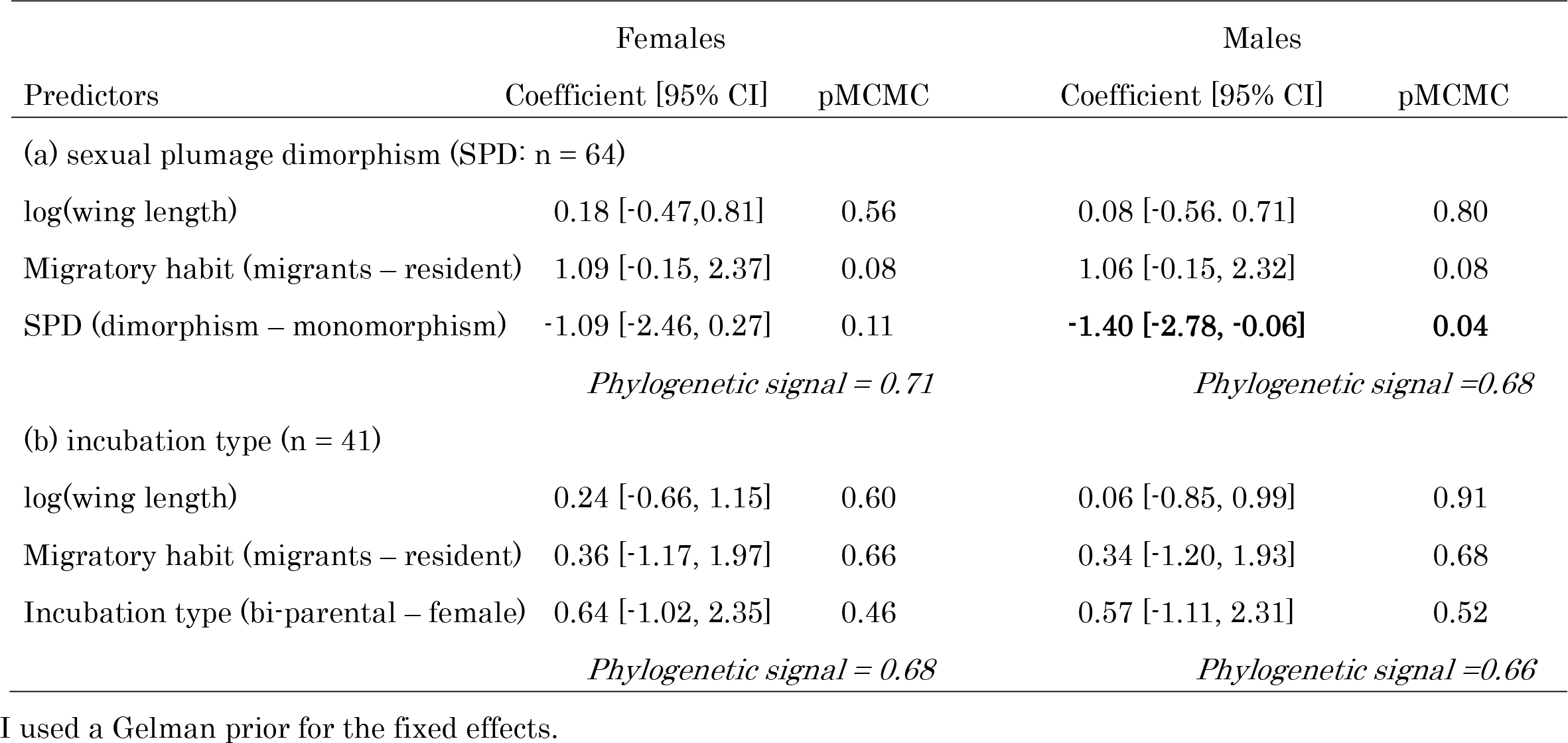
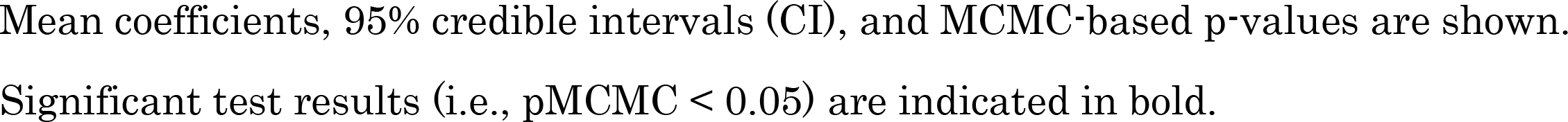
Multivariable Bayesian phylogenetic mixed model with an ordinal distribution predicting the degree of lore-forehead borderline (0: absent; 1: ambiguous; 2: obvious) in relation to measures of sexual selection (a: sexual plumage dimorphism and b: incubation type) and covariates in female and male hirundines after excluding species with pale lores.

|  | Females |  | Males |  |
| --- | --- | --- | --- | --- |
| Predictors | Coefficient [95% CI] | pMCMC | Coefficient [95% CI] | pMCMC |
| (a) sexual plumage dimorphism (SPD: n = 64) |  |  |  |  |
| log(wing length) | 0.18 [-0.47,0.81] | 0.56 | 0.08 [-0.56, 0.71] | 0.80 |
| Migratory habit (migrants – resident) | 1.09 [-0.15, 2.37] | 0.08 | 1.06 [-0.15, 2.32] | 0.08 |
| SPD (dimorphism – monomorphism) | -1.09 [-2.46, 0.27] | 0.11 | **-1.40 [-2.78, -0.06]** | **0.04** |
|  | *Phylogenetic signal = 0.71* |  | *Phylogenetic signal =0.68* |  |
| (b) incubation type (n = 41) |  |  |  |  |
| log(wing length) | 0.24 [-0.66, 1.15] | 0.60 | 0.06 [-0.85, 0.99] | 0.91 |
| Migratory habit (migrants – resident) | 0.36 [-1.17, 1.97] | 0.66 | 0.34 [-1.20, 1.93] | 0.68 |
| Incubation type (bi-parental – female) | 0.64 [-1.02, 2.35] | 0.46 | 0.57 [-1.11, 2.31] | 0.52 |
|  | *Phylogenetic signal = 0.68* |  | *Phylogenetic signal =0.66* |  |

I used a Gelman prior for the fixed effects.
Mean coefficients, 95% credible intervals (CI), and MCMC-based p-values are shown.
Significant test results (i.e., $p_{\text{MCMC}} < 0.05$ ) are indicated in bold.

**Figure S1.**
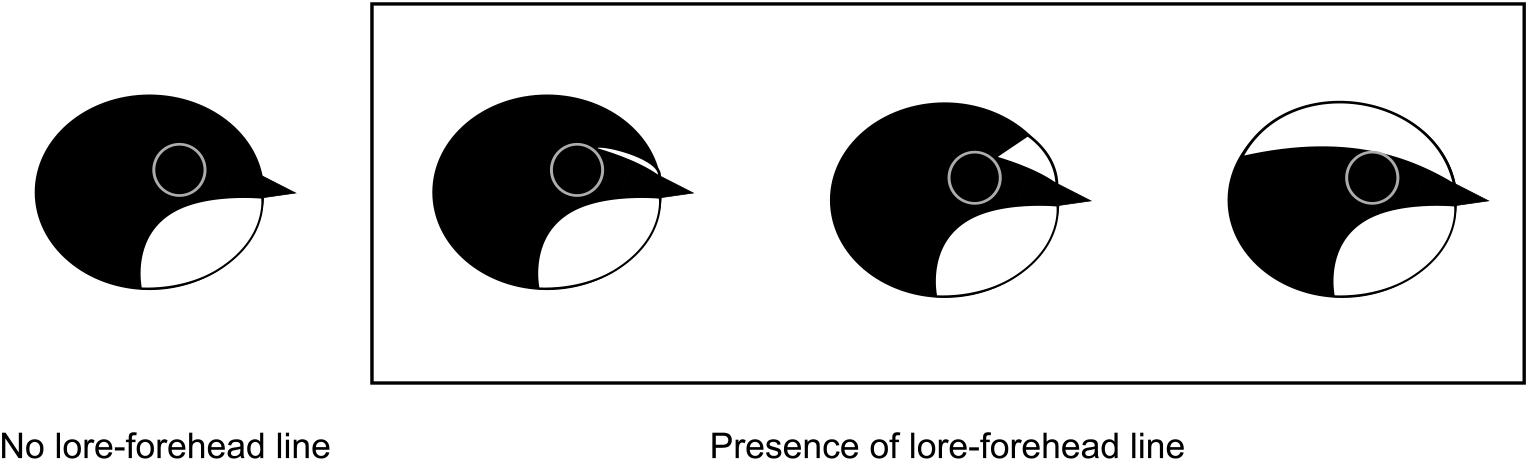
Lore-forehead borderline. Species with eyebrow stripes (including those with supraloral stripes), species with forehead patches, and species with caps were all regarded to have lore-forehead borderlines (note that presence/absence of throat patches was not used as a criterion to have lore-forehead borderline, though hirundines often have pale throat feathers). In other words, I focus solely on the contrasted “borderline” between lore and forehead regions.

**Figure S2.**
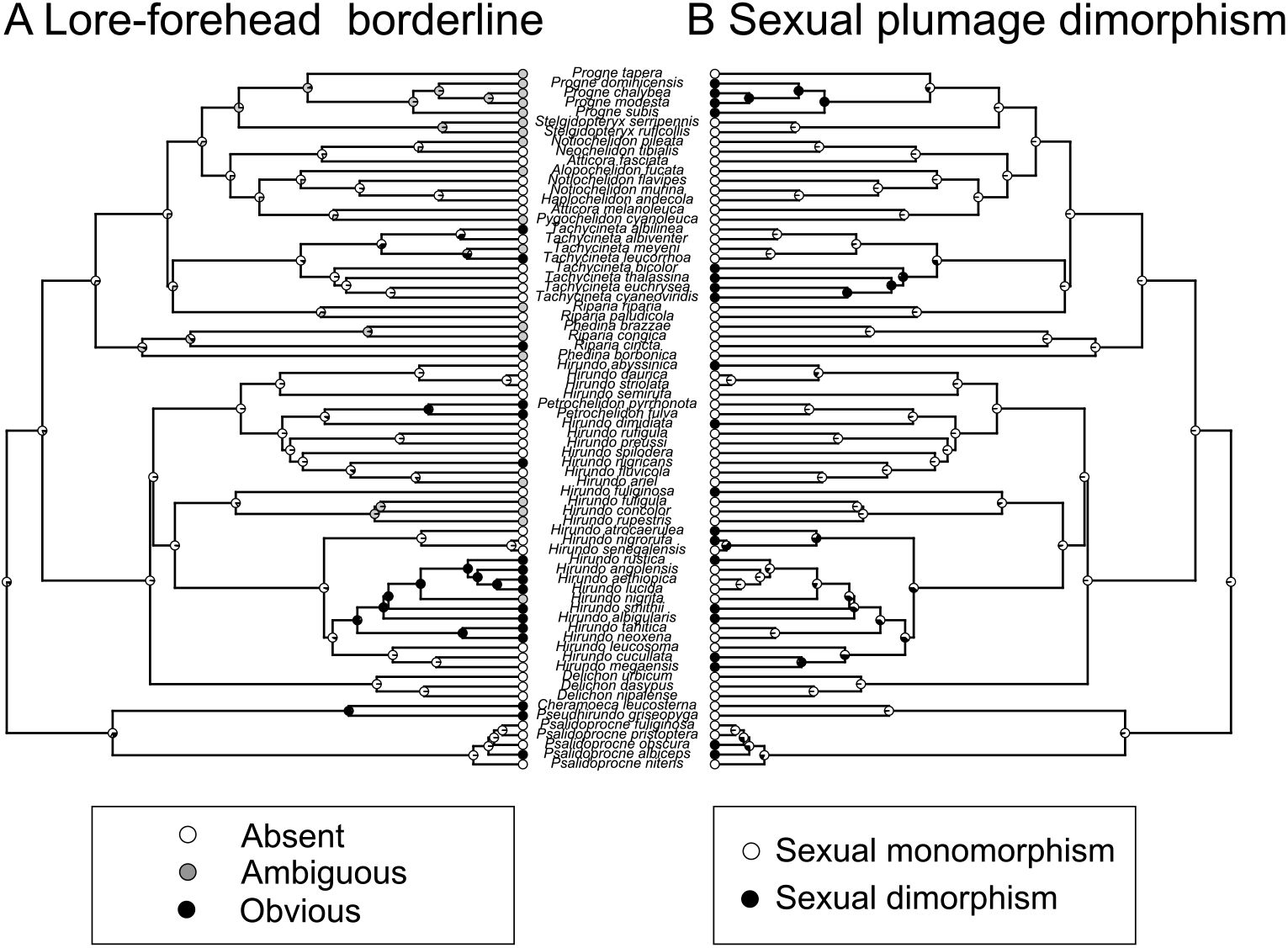
Examples of ancestral character reconstruction of the degree of lore-forehead borderline (left panel) and sexual plumage dimorphism (right panel) in swallows and martins (Aves: Hirundininae) using the functions ‘plotTree’ in the R package ‘phytools’ (Revell 2012). Black, grey, and white circles in tips in the left panel indicate species with obvious, ambiguous, and no lore-forehead borderline and black and white circles in tips in the right panel indicate species with sexually monomorphic and dimorphic plumage. Likewise, the proportions of black, grey, and white in nodes in the left panel indicate the probability of an ancestral state with obvious, ambiguous, and no lore-forehead borderline, and corresponding black and white proportion in the right panel indicate the probability of an ancestral state with sexual monomorphism and dimorphism.

**Figure S3.**
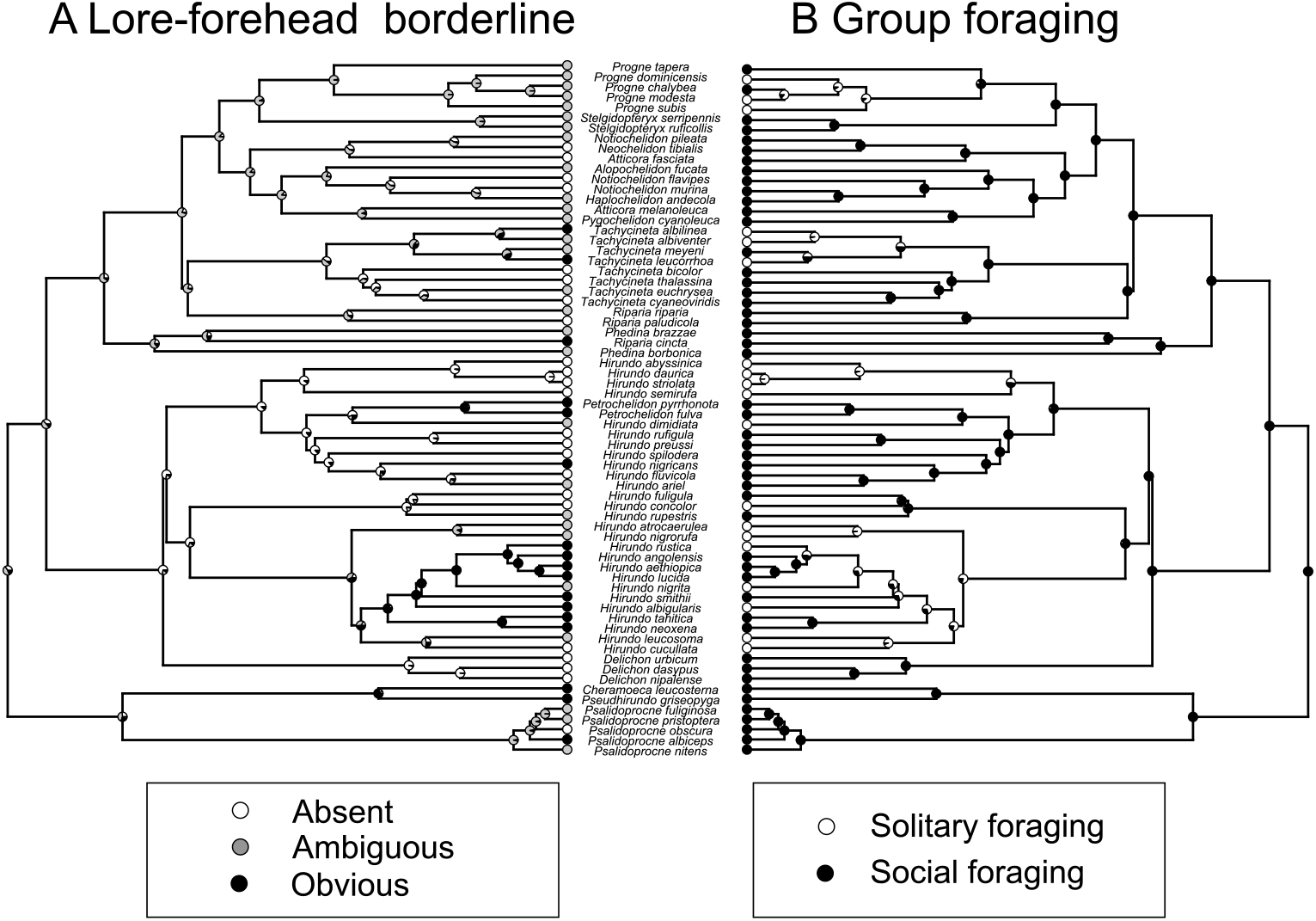
Examples of ancestral character reconstruction of the degree of lore-forehead borderline (left panel) and social foraging (right panel) in swallows and martins (Aves: Hirundininae) using the functions ‘plotTree’ in the R package ‘phytools’ (Revell 2012). Black, grey, and white circles in tips in the left panel indicate species with obvious, ambiguous, and no lore-forehead borderline and black and white circles in tips in the right panel indicate species with solitary and social foraging. Likewise, the proportions of black, grey, and white in nodes in the left panel indicate the probability of an ancestral state with obvious, ambiguous, and no lore-forehead borderline, and corresponding black and white proportion in the right panel indicate the probability of an ancestral state with solitary and social foraging.

**Figure S4.**
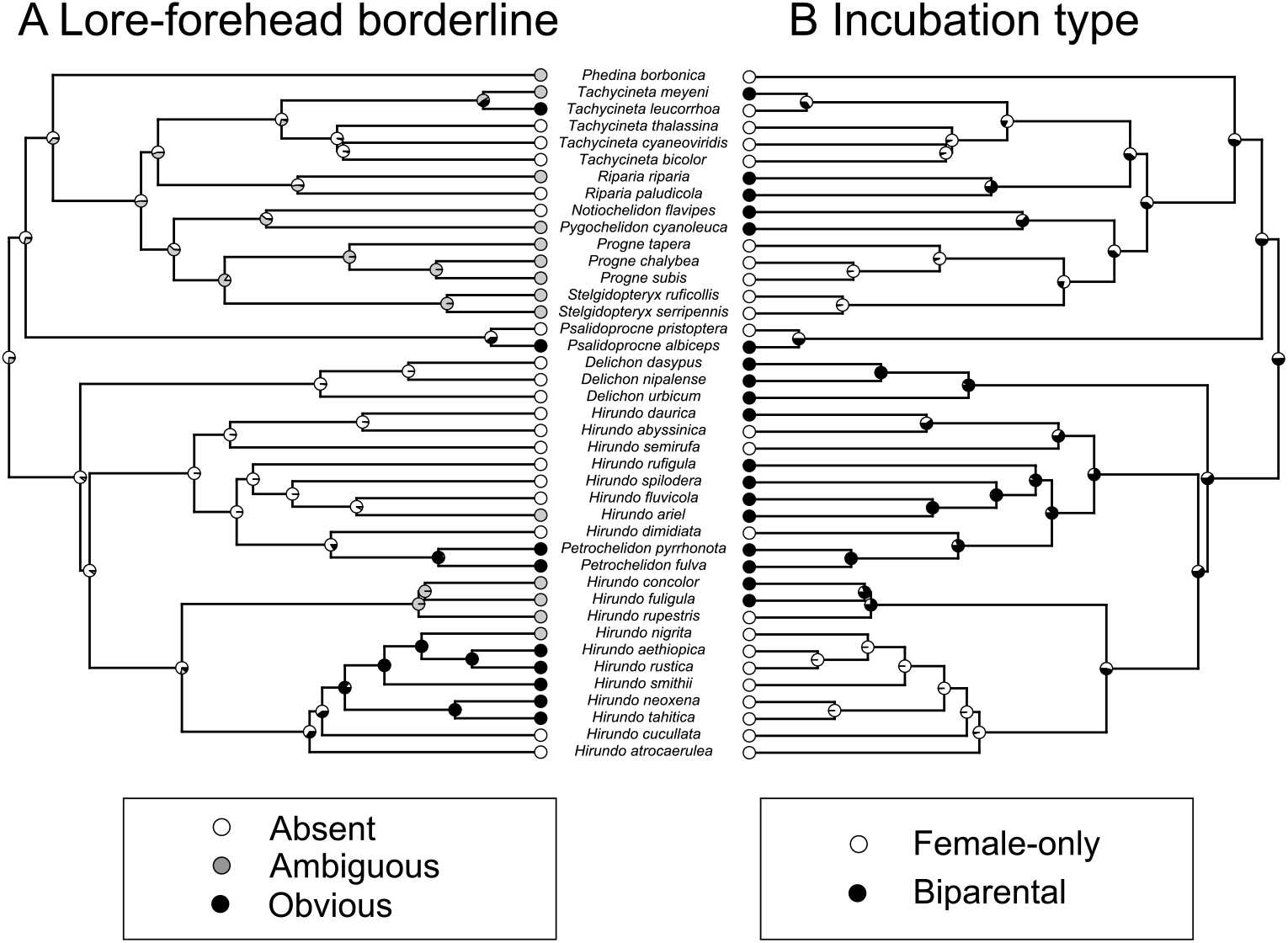
Examples of ancestral character reconstruction of the degree of lore-forehead borderline (left panel) and incubation type (right panel) in swallows and martins (Aves: Hirundininae) using the functions ‘plotTree’ in the R package ‘phytools’ (Revell 2012). Black, grey, and white circles in tips in the left panel indicate species with obvious, ambiguous, and no lore-forehead borderline and black and white circles in tips in the right panel indicate species with female-only and biparental incubation. Likewise, the proportions of black, grey, and white in nodes in the left panel indicate the probability of an ancestral state with obvious, ambiguous, and no lore-forehead borderline, and corresponding black and white proportion in the right panel indicate the probability of an ancestral state with female-only and biparental incubation.

**Figure S5.**
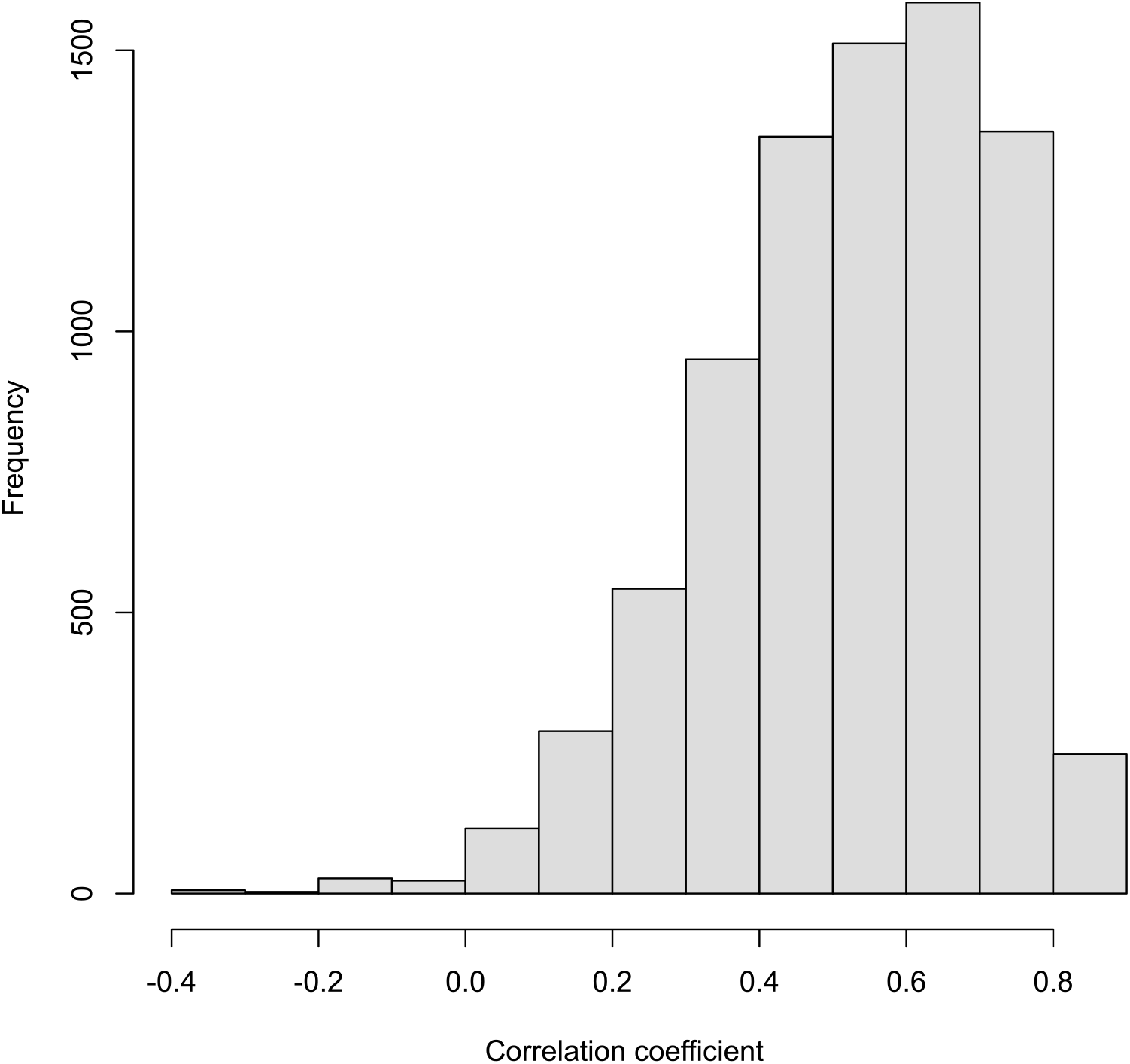
An example of the estimated posterior distribution of correlation coefficient between female lore-forehead borderline (absence = 0, presence =1) and prey size (small = 0, large = 1) in hirundines when applying function threshBayes in the R package phytools (with chains run for 1000000 generations, discarding the first 20% as burn-in). The mean correlation coefficient and P_MCMC_ values in this example are 0.53 and 0.01, respectively.

**Figure S6.**
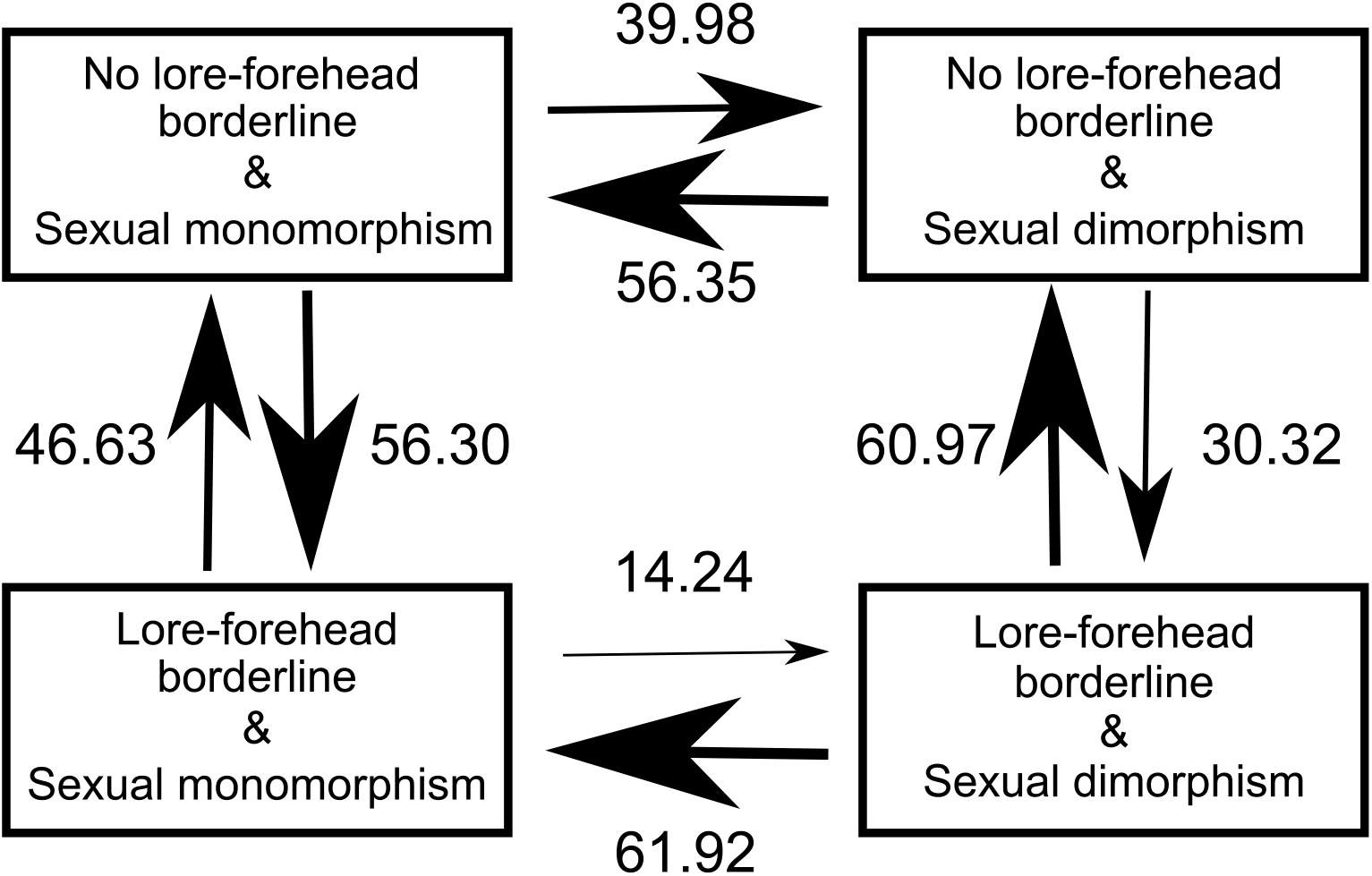
The most likely evolutionary transition between the presence and degree of male lore-forehead borderline in relation to sexual plumage monomorphism/dimorphism (see text) in the family Hirundinidae. Model-averaged transition rates, which are reflected by arrow size, are depicted.

**Figure S7.**
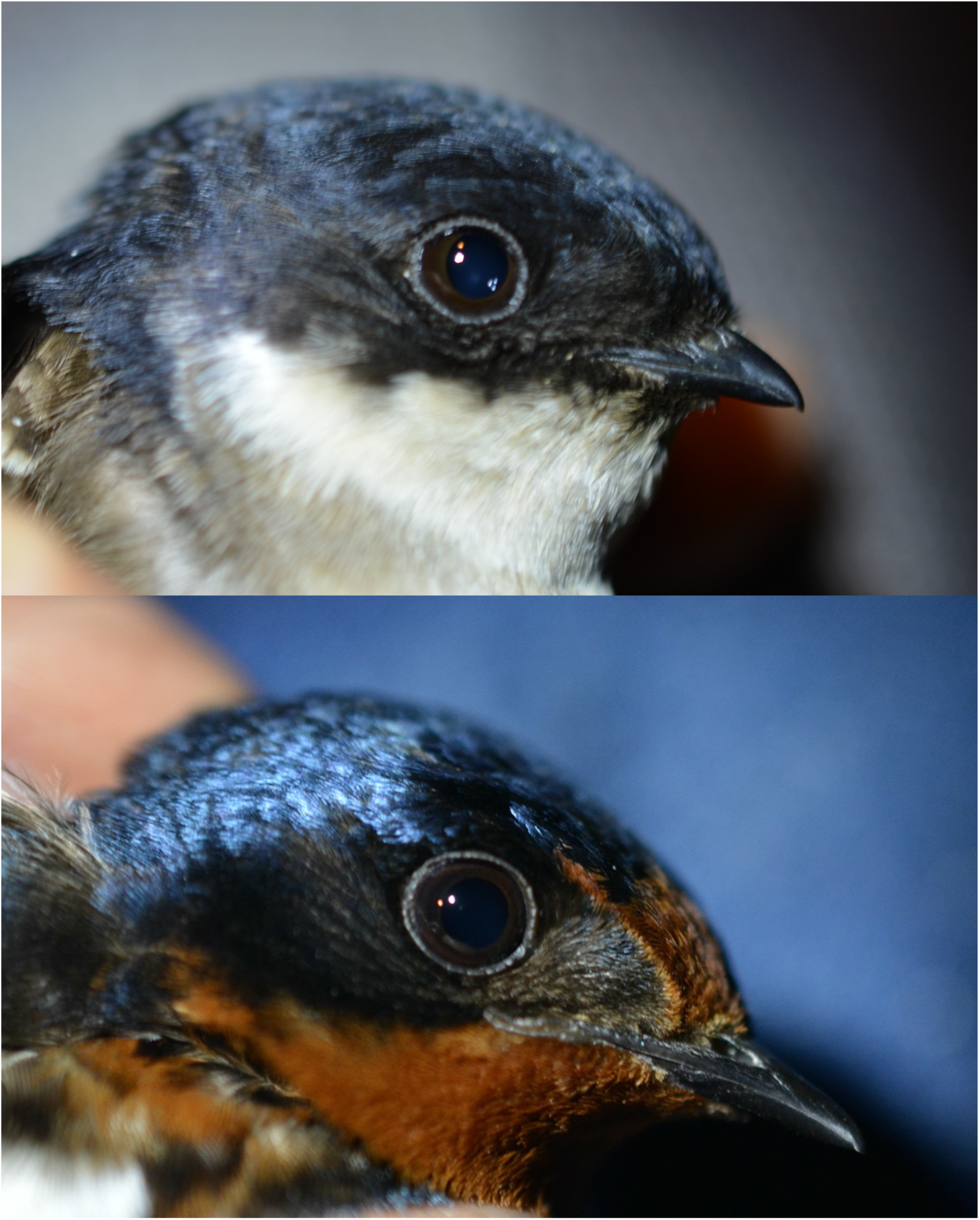
Closeup of the face of the Asian house martin (upper panel) and the barn swallow (lower panel). Note the clearcut lore-forehead borderline of the barn swallow.

